# Principles of ipsilateral and contralateral cortico-cortical connectivity in the mouse

**DOI:** 10.1101/033878

**Authors:** Alexandros Goulas, Harry BM Uylings, Claus C Hilgetag

**Author notes:** Corresponding author: Dr. Alexandros Goulas, Department of Computational Neuroscience, University Medical Center Hamburg-Eppendorf, Martinistr. 52, 20246 Hamburg, Germany.

## Abstract

Structural connectivity among cortical areas provides the substrate for information exchange in the brain and is characterized by the presence or absence of connections between specific areas. What principles govern this cortical wiring diagram? Here, we investigate the relation of physical distance and cytoarchitecture with the connectional architecture of the mouse cortex. Moreover, we examine the relation between patterns of ipsilateral and contralateral connections. Our analysis reveals a mirrored and attenuated organization of contralateral connections when compared to ipsilateral connections. Both spatial proximity and cytoarchitectonic similarity of cortical areas are related to the presence or absence of connections. Notably, our analysis demonstrated that these factors conjointly relate better to cortico-cortical connectivity than each factor in isolation, and that the two factors contribute differently to ipsilateral and contralateral connectivity. Distance is more tightly related to the presence or absence of ipsilateral connections, but its contribution greatly diminishes for contralateral connections, while the contribution of cytoarchitectonic similarity remains stable. Our results, conjointly with similar findings in the cat and macaque cortex, suggest that a common set of principles underlies the macroscale wiring of mammalian brains.

## Introduction

The connectional architecture of the mammalian cortex provides the anatomical substrate for the communication of its distinct elements. At the macro-scale level, such connectional architecture corresponds to the long-range white matter pathways linking the mosaic of areas of the cortical sheet (Sporns et al. 2005). Extensive invasive studies in animal models like the macaque monkey have uncovered a characteristic pattern of absence or presence of connections between specific cortical areas (e.g. Pandya and Yeterian 1990; Yeterian et al. 2012). However, few studies have aimed at uncovering the principles underlying the cortico-cortical connectional architecture in a systematic and quantitative way, an endeavor that is important for understanding the basic blueprint of the wiring of the cortex and identifying fundamental candidate neurodevelopmental mechanisms resulting in such wiring.

At least two wiring principles seem to underlie the cortico-cortical connectional architecture. The first principle is the physical distance between two cortical areas, that is, areas which are close to each other are likely to be connected while increasingly distant areas are less likely to be connected (e.g. Greilich 1984; Young 1992). This principle reflects a wiring cost reduction design (Ramón y Cajal 1899; Scannell et al. 1995; Kaiser and Hilgetag 2006). The second principle is grounded in the cytoarchitecture of cortical areas, suggesting a “similar prefers similar” wiring (Pandya and Yeterian 1990; Barbas 2015). Based on this principle, areas that are more similar in terms of their cytoarchitecture, for instance two agranular areas (areas lacking layer IV), are more likely to establish connections between them, while less similar areas, such as an agranular area and a granular area (an area possessing layer IV), are less likely to be connected. Both of the aforementoned principles have been shown quantitatively to relate to the presence or absence of connections between cortical areas of the cat (Beul et al. 2015a) and the macaque monkey (Beul et al. 2015b). Thus, cytoarchitectonic similarity and physical distance (hereafter simply referred to as distance) seem to constitute mammalian-general principles of cortico-cortical wiring. Further examination of the wiring of other mammalian brains is necessary to solidify such claim.

A limitation of most studies on cortico-cortical connectivity is the lack of a whole brain examination of ipsilateral and contralateral connections. Certain exceptions focusing on the macaque prefrontal cortex revealed a large overlap of the topography and high correlation of the strength of ipsilateral and contralateral connections, thus highlighting a mirrored organization of ipsilateral and contralateral connectivity (Barbas et al. 2005). Such insights foster the development of hypotheses on the relation of ipsilateral and contralateral connections in other mammalian species. Datasets offering information on contralateral connectivity also allow the examination of the relation of cytoarchitectonic similarity and distance to contralateral connections.

Recent efforts have generated valuable connectivity data of the mouse cortex (Oh et al. 2014; Zingg et al. 2014). These datasets constitute the current best estimate of cortico-cortical connectivity in the mouse. In the current study we used these wiring diagrams and adopted a quantitative approach to examine the relation of ipsilateral and contralateral connections in the mouse cortex and the relation of connectivity to the spatial proximity and cytoarchitectonic similarity of cortical areas.

## Materials and Methods

### Connectivity data

The dataset used for our main analysis is the Allen Mouse Connectivity Atlas (http://connectivity.brain-map.org/). The mouse wiring diagram was mapped by employing the recombinant adeno-associated virus expressing enhanced green fluorescent protein as an anterograde tracer. For constructing a large scale connectivity map, the Allen Reference Atlas (http://mouse.brain-map.org/static/atlas) was used. In total 295 non-overlapping structures (cortical areas, subcortical nuclei etc.) were taken into account and the majority of the injections involved distinct structures. Altogether 469 injected brains of C57BL/6J male mice were included in the construction of the large-scale connectivity matrix through constrained optimization. The constrained optimization set about two thirds of all possible connections to zero (absent). P-values were estimated for the remaining non-zero weights with linear regression (see Oh et al. 2014 for details on the estimation of the p-values). This procedure resulted in connectivity matrices involving 213 structures. Details on the informatics pipeline, quality controls and estimation of the inter-areal connectivity matrix are provided in (Oh et al. 2014). The connectivity matrices from (Oh et al. 2014) were obtained from the Open Connectome project (http://www.openconnectomeproject.org/). The .graphml file was converted to .gml with the online Open Connectome project conversion tools (http://mrbrain.cs.jhu.edu/graph-services/convert/)· Lastly, the data from the .gml file were imported in Matlab and converted into a directed graph with the aid of Matlab scripts (http://www.mathworks.de/matlabcentral/fileexchange/45741-read-gml). In the current study we focused only on cortico-cortical connections involving the 38 cortical areas of the Allen Mouse Connectivity Atlas (Fig. 1). Connections were considered present if they exhibited a p-value, obtained from the linear regression, below 0.05 and all remaining connections were considered absent. All connections were treated as binary unless otherwise stated, that is, when the strength of the connections was examined. We used as connectivity strength the so-called normalized connectivity strength that quantifies the amount of signal detected in a target area after infecting one voxel in the source area (see Oh et al. 2014 for details). Moreover, for assessing if the results are driven primarily by homotopic connections (connections linking the same area in the two hemispheres), we performed the analysis with and without the homotopic connections. The results reported below are derived from the analysis without the homotopic connections (except from the homotopic strength analysis (see next paragraph)). Inclusion of homotopic connections did not change the results.

**Figure 1.**
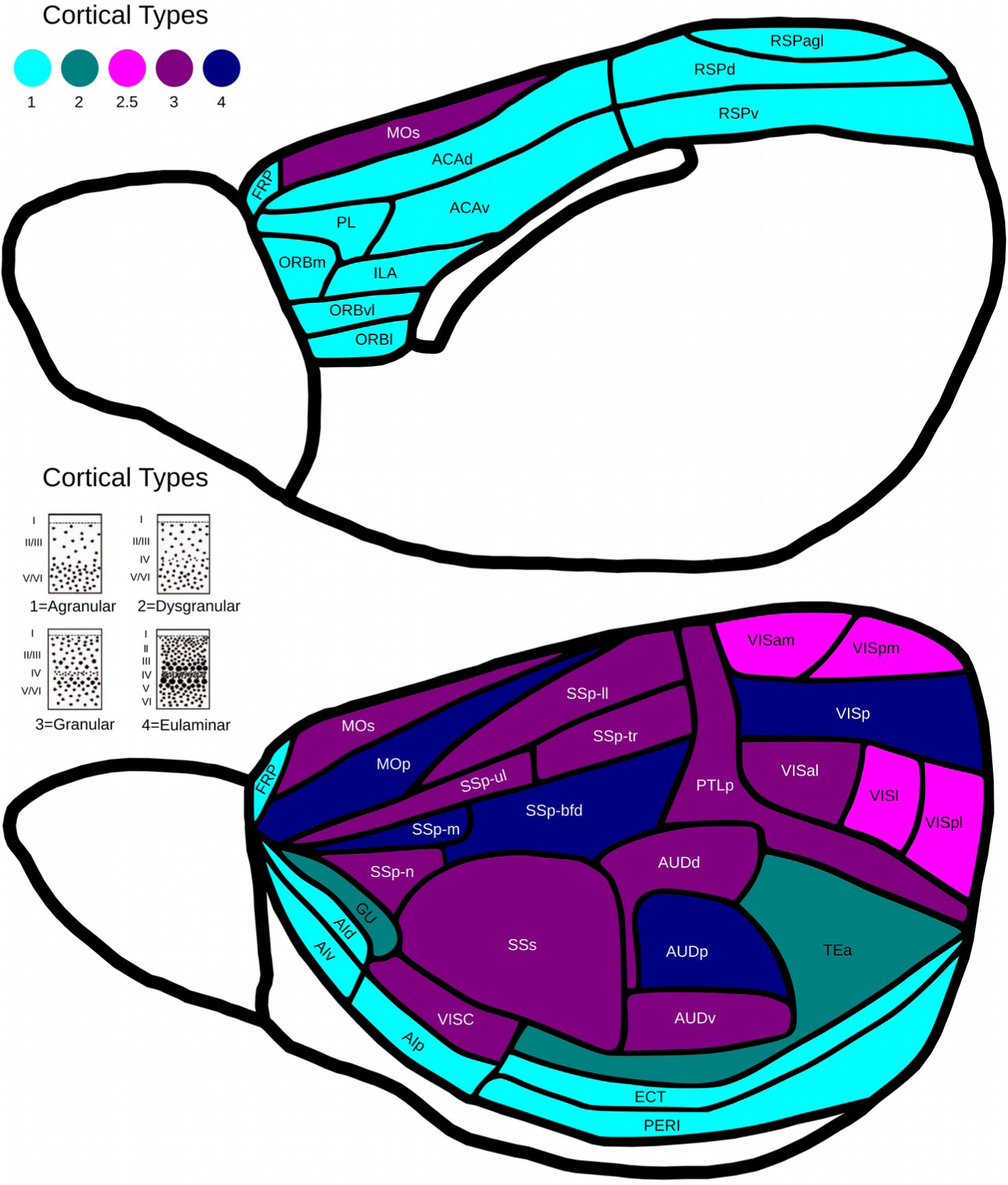
Schematic depiction of the cortical areas of the mouse cortex based on the Allen Reference Atlas. Note that this drawing is an approximation for offering an overview of the cortical areas in the mouse. Colors denote the distinct cortical types as an ordinal scale, with 1 denoting the less eulaminated areas and 4 the more eulaminated. An intermediate type (2.5) denotes areas that exhibit substantial within-area heterogeneity, that is, a combination of cortical type 2 and 3. The actual Allen Reference Atlas is available online (http://mouse.brain-map.org/static/atlas). For abbreviations of the cortical areas see (Oh et al. 2014). For a discussion on the nomenclature and other parcellation schemes of the mouse cortex see (Van De Werd and Uylings 2014) and (Paxinos and Franklin 2013).

### Relation of ipsilateral and contralateral connections

Previously Oh et al. (Oh et al. 2014) used simple metrics for comparing the ipsilateral and contralateral connections, that is, the correlation and ratio of ipsilateral and contralateral connection strengths at the whole brain level. Here we employ such metrics, as well as additional topological metrics, conjointly with statistical inference for examining the similarity of ipsilateral and contralateral cortico-cortical connections. We adopt a similar approach as previously done for the macaque prefrontal cortex (Barbas et al. 2005). Specifically, the following metrics were computed. For assessing the overall topological similarity of the ipsilateral and contralateral matrices we used the edit distance (e.g. Trusina et al. 2005) that assesses the number of insertion/deletion operations needed to convert one matrix to the other. This analysis takes into account only the topology, that is, the presence or absence of connections. In order to assess the relation of the strength of ipsilateral and contralateral connections, we computed the Spearman’s rank correlation between the two matrices. In addition to this global analysis, we performed an area-wise analysis by computing the following metrics. First, for each area the ratio of the sum of the strength of ipsilateral versus contralateral connections was computed (Barbas et al. 2005). Second, the number of areas common to the ipsilateral and contralateral projection patterns of an area was assessed with the Jaccard index, defined as the intersection versus the union of ipsilaterally and contralaterally connected areas. Third, we estimated the similarity between the ipsilateral and contralateral connectivity patterns by taking into account the weights of the connections and calculating Spearman’s rank correlation between the contralateral and ipsilateral connectivity profiles. The null values for these metrics were computed from 1000 null models for the ipsilateral and contralateral connectivity matrices, matched in node (number of areas), edge (number of total connections) and degree distribution (number of connections of an area) (Rao and Bandyopadhyay 1996). Lastly, we assessed if connections between homotopic areas (e.g. the connection between the frontopolar cortex in the left and right hemisphere) were significantly stronger than the rest of the contralateral connections.

### Relating cytoarchitectonic similarity and distance to cortico-cortical connections

For evaluating the role of distance, we used the Euclidean distance between the center of mass of the cortical areas. This distance functions as a proxy of the length, and consequently wiring cost, of the connection between two areas. Euclidean distances between the center of masses of the cortical areas are provided in (Oh et al. 2014). We expected that areas separated by short Euclidean distances are more frequently connected while increasingly distant areas are less frequently connected. For examining the relation between presence or absence of connectivity and cytoarchitectonic similarity, we classified the 38 areas in 4 cortical types: 1=agranular (absence of layer IV), 2=dysgranular (layer IV is not easily discernible), 3=granular (presence of layer IV) and 4=eulaminar (more distinct differentiation of layers accompanied by a more prevalent layer IV) (Fig. 1). To this end, high resolution Nissl stained sections were used from a previous cytoarchitectonic study in the mouse (Van De Werd and Uylings 2014) in conjunction with the Paxinos and Franklin’s Mouse Atlas (Paxinos and Franklin 2013). It should be noted that such an ordinal scale of cortical types has been shown to exhibit a high positive correlation with objective measures of cytoarchitecture like neuronal density (Dombrowski et al. 2001) thus capturing essential cytoarchitectonic differences of cortical areas. Since the wiring of the cat and macaque cortex obeys a “similar prefers similar” principle (Beul et al. 2015a; Beul et al. 2015b; Pandya and Yeterian 1990), we expected that mouse cortico-cortical connections are more likely to be present between areas of similar cortical types. In order to express the cytoarchitectonic difference of two areas, the index |Δ| was used denoting the absolute difference between the cortical type of a pair of areas (e.g. Barbas et al. 2005). Fig. 2 offers a visual summary of the overall approach.

**Figure 2.**
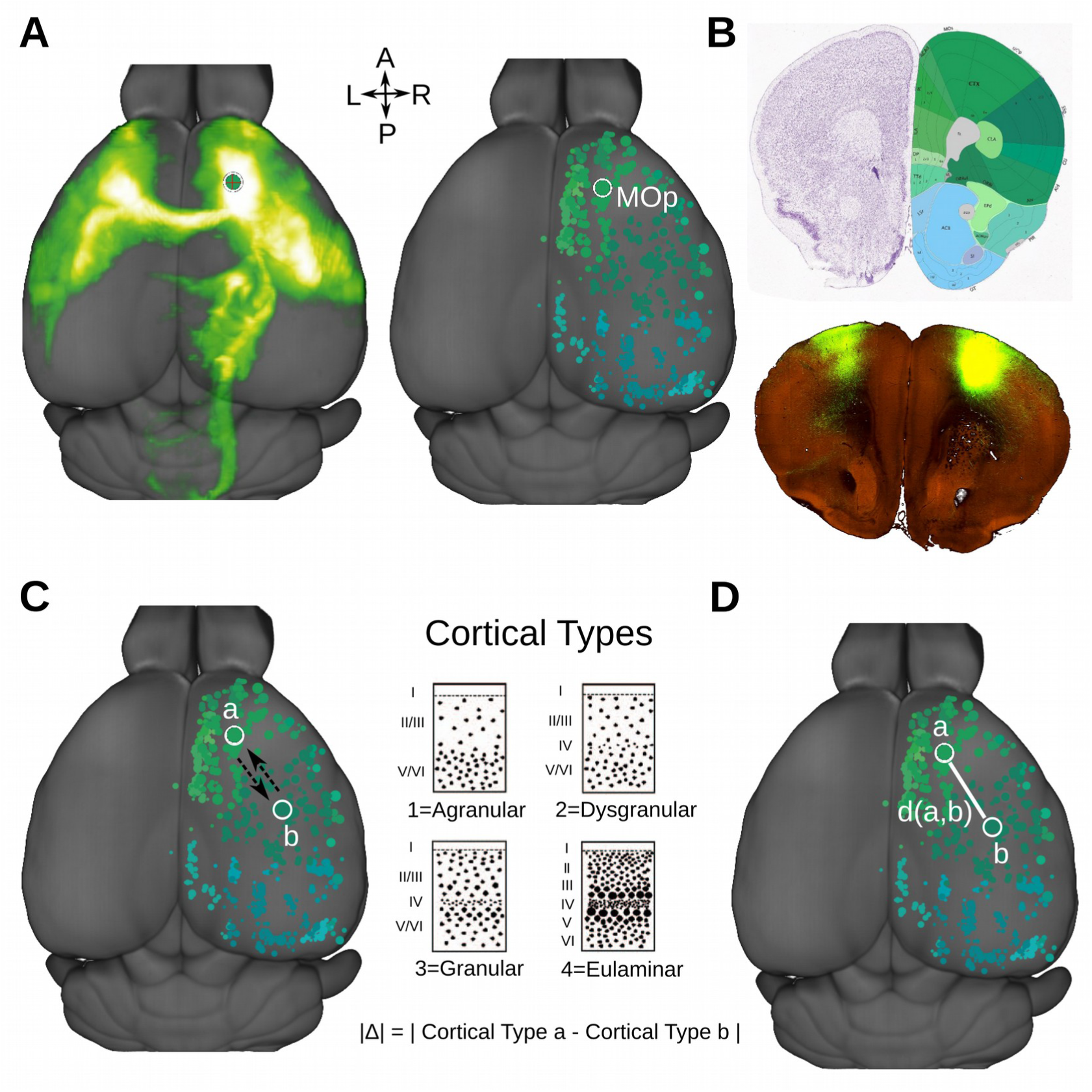
Example of data used and schematic depiction of the factors related to presence or absence of connectivity **A**. Ipsilateral and contralateral connectivity pattern of the primary motor area (MOp). **B**. Axial slice depicting the pattern of connectivity of area MOp alongside with a Nissl stained section and the corresponding parcellation based on the Allen Reference Atlas. **C**. Cytoarchitectonic similarity of cortical areas was estimated by an index |Δ| computed for each pair of areas as the absolute difference of their cortical type. Note that this index is computed for a pair of areas irrespective of the presence or absence of connection(s) (depicted with dashed lines) in between them. **D**. The wiring cost of a potential connection between two areas was approximated by the physical distance between the two areas, i.e. the Euclidean distance between their centers of mass.

For illustrating the relation of distance and cytoarchitectonic similarity to presence or absence of connections in an intuitive manner, we binned the Euclidean distances and the |Δ| values. For each bin the number of present connections was expressed as a proportion of present connections divided by the number of possible connections for the current bin.

We used nominal logistic regression for examining the relation between presence or absence of connections (dependent variable) and distance and cytoarchitectonic similarity (predictors). The two predictors range in different units and thus were normalized to the 0–1 interval. An initial analysis examined all connections (ipsilateral and contralateral) simultaneously and used all predictors. For this analysis an additional categorical predictor coding for the contralateral (=1) and ipsilateral (=0) connections was used. This analysis also examined the interactions of this categorical predictor with the predictors of distance and cytoarchitectonic similarity. A subsequent analysis was run on the ipsilateral and contralateral connections separately. In this step, we examined the contribution of distance and cytoarchitectonic similarity separately and then conjointly for assessing their unique contribution. The quality of the logistic regression models were assessed with McFadden’s pseudo-R^2^, which quantifies the improvement of the likelihood of the model when compared to a null model (a model containing only the intercept term). The statistical difference of the likelihood of the bivariate and univariate models was assessed with the likelihood ratio test (e.g. Vidakovic 2011). The likelihood ratio test explicitly addresses if the addition of a predictor significantly improves a model, thus assessing if a more “complex” model (with more predictors) should be favored.

In order to assess the generalizability of models build with each predictor in isolation or conjointly, a prediction analysis was conducted. To this end, the coefficients from the nominal logistic regression were used to predict the presence or absence of connections with a model build with distance, cytoarchitectonic similarity, or both as predictors. The quality of such predictions was assessed by computing receiver operating characteristic (ROC) curves and the corresponding areas under curve (AUC) for the original and null (with shuffled labels) predictions. It should be noted that in a prediction analysis the addition of a predictor does not necessarily lead to higher AUC values. Thus, a model leading to statistically significant higher AUC values should be favored. The predictions were computed 100 times each time using 80% of the available data to build the model (drawing with replacement) and the rest of the data serving as a test set. The percentage of data used (varying from 80% to 90%) did not change the results. The prediction analysis was carried out separately for the ipsilateral and contralateral connections.

### Control analyses

A series of control analyses was performed to ensure robustness of the results and their independence from parameters in the analysis.

First, an additional independent dataset on male C57BL/6J mouse cortico-cortical connectivity was tested (Zingg et al. 2014) (See Supplementary Data).

Second, for the connectivity matrices of the main analysis obtained from the Allen Mouse Connectivity Atlas, a more conservative p-value threshold was used, that is connections with a p-value smaller than 0.01 were considered present and all other connections were considered absent.

Third, all connectivity datasets were subject to the following control analyses: (1) Two areas were randomly excluded and all analyses were redone for these reduced cortico-cortical matrices. This control analysis investigated if the choice of examining a particular set of cortical areas has an impact on the results. (2) 10% of connections were assumed to be present, and in a separate analysis absent, and all the analyses were redone for these “randomly enriched” (or “randomly reduced”) matrices. Thus, this simple scenario simulates a situation were connections are wrongly assumed as absent or present due to potential biases in the pipeline resulting in the connectivity matrices. (3) 10% of the cortical areas were assigned to a different cortical type by randomly moving them to a category denoting an immediately higher or lower cortical type. For instance, area MOs could potentially be assigned to cortical type 2 to reflect evidence for a less eulaminated medial part of MOs (Van De Werd et al. 2010). This control analysis addressed if small changes in our observer-dependent qualitative assignments impact the results.

A last control analysis aimed at taking into account the functional modality similarity of cortical areas. Previous findings at the microscopic connectivity level, that is pyramidal cell-to-pyramidal cell connectivity in the mouse primary visual cortex, suggests that cells are more probably connected if they have similar functional profiles (similar orientation preference of visual stimuli) (Ko et al. 2011). At the macroscale level, cortico-cortical connections in the cat cortex have been suggested to obey a functional similarity rule (Lee and Winer 2008a). Therefore, we tested if functional modality similarity of the mouse cortical areas “explains away” the relation between the presence or absence of connections and cytoarchitectonic similarity. To this end, we grouped the cortical areas in functional modalities and used an additional predictor in the nominal logistic regression coding for the functional modality similarity of two cortical areas (see Supplementary Data). All analyses were conducted in Matlab (8.1.0.604 R2013a) (Mathworks).

## Results

The results are structured by first presenting the ipsilateral and contralateral topological results and second the relation of distance and cytoarchitectonic similarity with presence or of connections.

### Relation of ipsilateral and contralateral connectivity patterns

Examination of the binary ipsilateral and contralateral connectivity matrices revealed that they are more similar than expected by chance (Edit Distance_Original_ 0.32 Edit Distance_null_ mean=0.71 std=0.02, p<0.001). Furthermore, a conserved connectivity strength pattern, that is, correlation of strength of ipsilateral and contralateral connections, was observed (rho=0.63, p<0.001). The area-wise analysis revealed that the ipsi/contra ratio of the strength of the areas was much higher than 1, on average 6.14, indicating much stronger ipsilateral connections (Fig. 3 A). A striking “outlier” was the ~35 times more prominent strength for ipsilateral versus contralateral connections for two areas (Gu and VISal). Despite the large differences in strength, the overall pattern of ipsilateral and contralateral connections was moderately to highly correlated, indicating a relatively high similarity of the ipsilateral and contralateral connectivity patterns for the majority of cortical areas (Fig. 3 B). Moreover, there was a moderate to high proportion of common areas in the ipsilateral and contralateral connectivity profile of each area (Fig. 3 C). The rendering of the ipsi/contra metrics across the cortical sheet (Fig. 3) indicates no systematic prominence of these metrics to specific sets of cortical areas or lobes as suggested for the human brain (Wang et al 2014). Hence, it appears that there is no clear segregation of cortical areas based on the ipsi/contra metrics (see also Discussion). Conjointly, the above results indicate that as a whole the topology and strength of ipsilateral and contralateral connection patterns are more similar than expected by chance, despite that contralateral connections are weaker than ipsilateral ones. Hence, the contralateral connectivity pattern appears to be an attenuated mirrored version of the ipsilateral pattern. Lastly, the homotopic connections were significantly stronger than the rest of contralateral connections (by a factor of x2.6 compared to contralateral non-homotopic connections, p<0.001, permutation test). This difference was also significant if distance was taken into account. Importantly, the above analysis involving the strength of connections led to the same qualitative results if the strength of connections were converted to “connectivity strength”, “connectivity density” or “normalized connectivity density” (see (Oh et al. 2014) for details).

**Figure 3.**
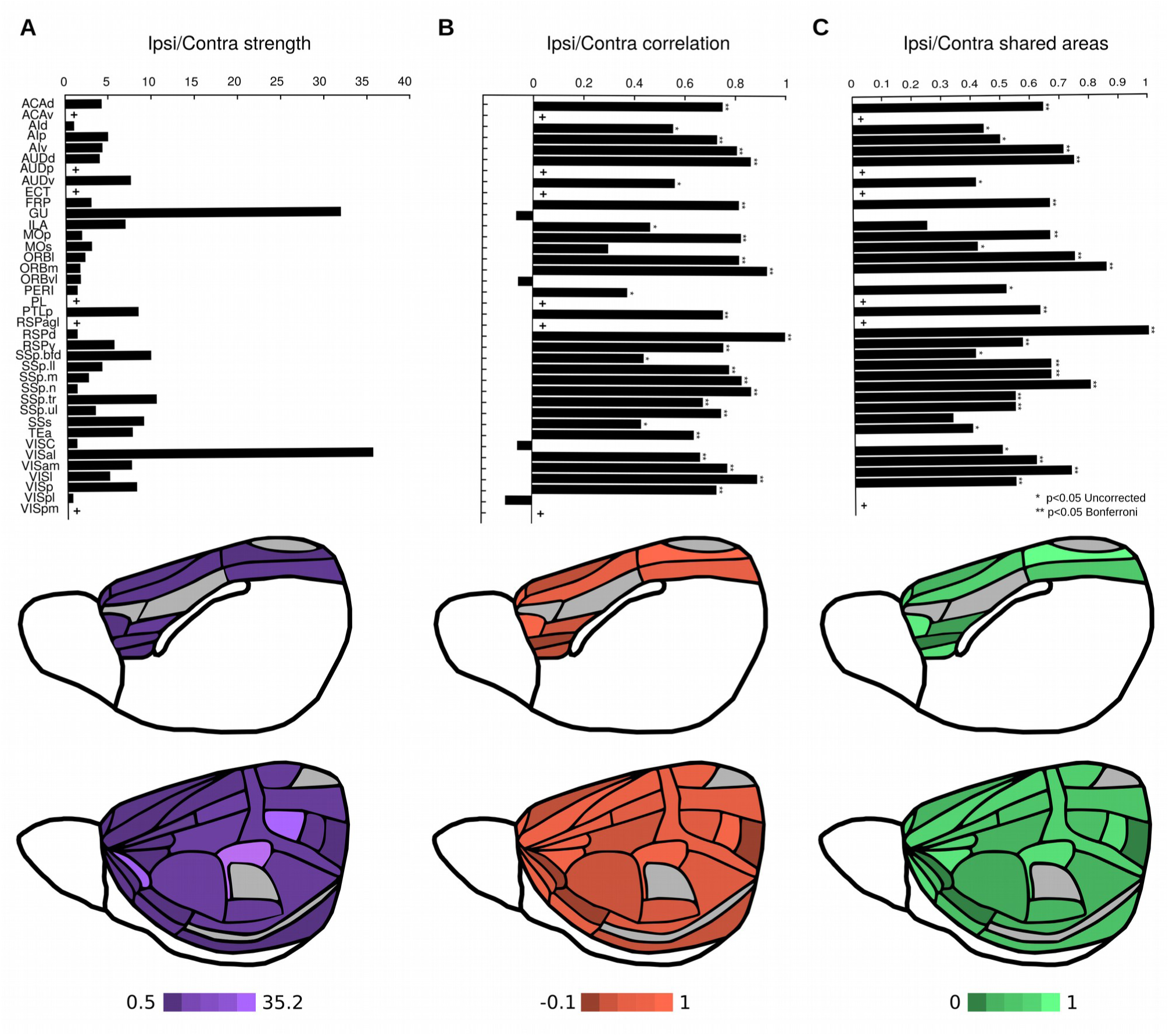
Ipsilateral and contralateral connectivity patterns. **A**. Ratio of ipsilateral over contralateral connection strength for each area. **B**. Correlation of the ipsilateral and contralateral connectivity profile for each area. **C**. Amount of shared areas, i.e. the Jaccard index of areas that are part of the ipsilateral and areas that are part of the contralateral connectivity profile. Based on the aforementioned metrics, no clear segregation of areas suggesting dichotomies such as primary versus non-primary is evident. The asterisks denote the significance of each area-wise metric in panels B and C. Significance was established via comparisons with metrics derived from null models (see Materials and Methods). Note that the areas marked with + do not have contralateral connections considered as “present” in the current p-value threshold (p<0.05).

### Relation of physical distance and cytoachitecture with presence or absence of connections

Increasing distance of areas and increasing cytoarchitectonic difference of cortical areas was accompanied by the occurrence of fewer connections between them. This relation was observed both for ipsilateral and contralateral connections (Fig. 4).

**Figure 4.**
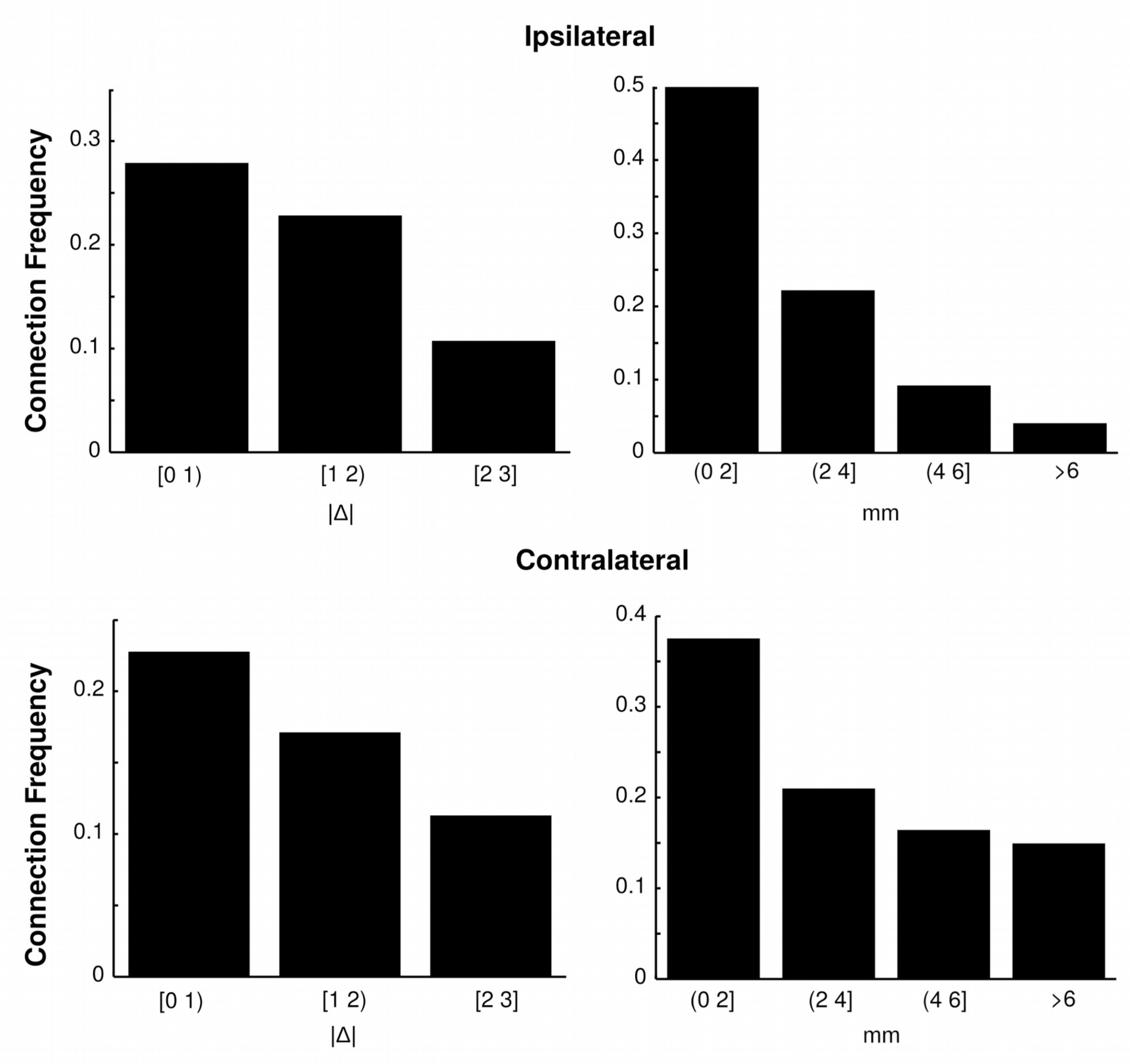
Frequency of present connections for different ranges of physical distance and cortical type differences. The connection frequency was estimated by dividing the number of existing connections by the number of possible connections for each interval of cytoarchitectonic similarity and distance. Note that for both ipsilateral and contralateral cases, increasing physical distance of areas and increasing cytoarchitectonic differences of areas were accompanied by a less frequent number of connections.

The nominal logistic regression model that took into account simultaneously all connections (ipsilateral and contralateral) revealed a significant contribution of distance (beta=-5.33 p<0.001) and cytoarchitectonic similarity (beta=-1.30 p<0.001). The negative sign of the regression coefficients indicates that an increase in distance or cytoarchitectonic difference is accompanied by a decrease in the probability of a connection being present compared to the probability of a connection being absent. In addition, the categorical predictor coding for the ipsilateral and contralateral connections was significant (beta=-1.61 p<0.001). The negative sign of the regression coefficient for this predictor reflects that the probability of a connection being present compared to the probability of a connection being absent decreases when shifting from the ipsilateral to the contralateral category. Moreover, the interaction between cytoarchitectonic similarity and the ipsilateral/contralateral categorical predictor was not significant (beta=0.10 p>0.1). This result indicates that the influence of cytoarchitectonic similarity on the presence or absence of connections does not depend on whether a connection belongs to the ipsilateral or contralateral category. On the other hand, the interaction between distance and the ipsilateral/contralateral categorical predictor was significant (beta=4.57 p<0.001). This result indicates that the influence of distance on the presence or absence of connections depends on whether a connection belongs to the ipsilateral or contralateral category.

### Relation of distance and cytoachitectonic similarity with ipsilateral and contralateral connections

We subsequently examined the contralateral and ipsilateral connections separately. It should be noted that the two predictors were almost orthogonal (rho=0.11 for ipsilateral connections, rho=0.01 for contralateral connections). Examination of each predictor in isolation, revealed that both distance and cytoachitectonic similarity were significantly related to the presence or absence of ipsilateral and contralateral connections (univariate model Table 1). The conjoint examination of physical distance and cytoachitectonic similarity again highlighted both predictors as significantly related to presence or absence of connections (bivariate model Table 1). The bivariate model was significantly better than the univariate models for the ipsilateral connections (Likelihood ratio test: 184.3 32.6 (p<0.001) when comparing the log likelihood of the bivariate model and the model built only on cytoachitectonic similarity and distance respectively). The bivariate model was also significantly better than the univariate models for the contralateral connections (Likelihood ratio test: 8.6 (p<0.05) 29.3 (p<0.001) when comparing the log likelihood of the bivariate model and the model built only on cytoachitectonic similarity and distance respectively).

**Table 1.** Results of the nominal logistic regression for the univariate and bivariate model for the data from the Allen Mouse Connectome Atlas. Note that the coefficients have a negative sign indicating that a unit increase in cortical type or physical distance leads to a decrease of the probability of a connection being present compared to the probability of a connection being absent (absent connections functioned as the “reference category” for the logistic regression). McFadden’s pseudo R^2^ ranges from 0 to 1 with higher values indicating a better model (increased improvement in terms of the likelihood of the model when compared to the null model, that is, a model only with the intercept term). The McFadden’s pseudo R^2^ values indicate a good model for the ipsilateral connections and a rather poor fit for the contralateral connections.

|  | Ipsilateral Univariate Model |  |  |  |  |
| --- | --- | --- | --- | --- | --- |
|  | Beta | 95% CI |  | p-value | McFadden's pseudo R<sup>2</sup> |
| Cytoarchitecture | -1.58 | -1.80 | -1.35 | 0.001 | 0.04 |
| Physical Distance | -4.86 | -5.24 | -4.47 | 0.001 | 0.15 |
|  | Contralateral Univariate Model |  |  |  |  |
| Cytoarchitecture | -1.23 | -1.46 | -1.01 | 0.001 | 0.02 |
| Physical Distance | -0.97 | -1.27 | -0.66 | 0.002 | 0.01 |
|  | Ipsilateral Bivariate Model |  |  |  |  |
| Cytoarchitecture | -1.30 | -1.76 | -0.84 | 0.001 | 0.17 |
| Physical Distance | -4.59 | -5.35 | -3.84 | 0.001 |  |
|  | Contralateral Bivariate Model |  |  |  |  |
| Cytoarchitecture | -1.20 | -1.68 | -0.78 | 0.001 | 0.03 |
| Physical Distance | -0.90 | -1.57 | -0.37 | 0.003 |  |

The prediction analysis results were as follows. For the ipsilateral connections, using only distance as predictor led to the highest AUC=0.76 (p<0.01, permutation test). Using only cytoachitectonic similarity as a predictor led to an AUC=0.64 (p<0.01, permutation test). The use of both predictors led to slightly better predictions (AUC=0.78, p<0.01, permutation test) compared to the ones using each predictor separately (Fig. 5). Comparing the AUC for the ipsilateral predictions revealed that significantly better predictions are obtained with distance when compared to the ones using cytoachitectonic similarity. Moreover, the combination of distance and cytoachitectonic similarity led to significantly better predictions compared to the ones obtained from using each predictor separately (p<0.001, permutation tests). For the contralateral connections, the highest AUC was observed for cytoachitectonic similarity (AUC=0.61 p<0.01, permutation test). Distance led to an AUC=0.55 (p<0.01, permutation test). The use of both predictors led to slightly better predictions (AUC=0.63, p<0.01, permutation test) compared to the ones using each predictor separately (Fig. 5). Comparing the AUC for the contralateral predictions revealed that significantly better predictions were obtained with cytoachitectonic similarity when compared to the ones using distance. Moreover, the combination of distance and cytoachitectonic similarity led to significantly better predictions compared to the ones obtained from using each predictor separately (p<0.001, permutation tests).

**Figure 5.**
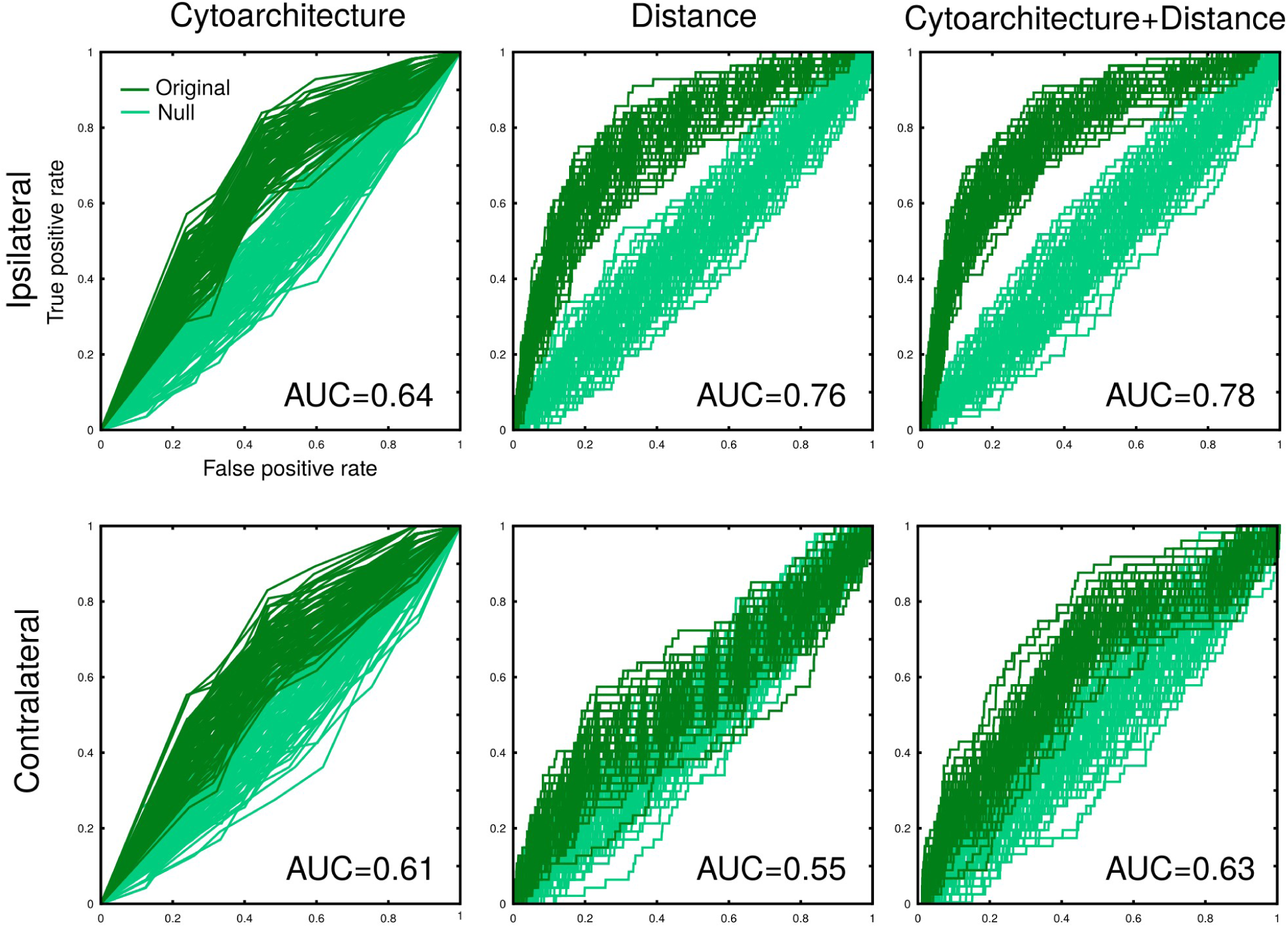
Connectivity prediction analysis based on cytoarchitectonic similarity and physical distance. For each analysis, 100 predictions and corresponding ROC curves were constructed by drawing with replacement. ROC curves are constructed for the original predictions with the true labels (present versus absent connection) as well as for null predictions obtained with shuffled labels. The quality of the prediction is quantified with the AUC. The depicted AUC values are the mean AUC values for the original predictions across 100 iterations. All AUC values are significantly different from the AUC values obtained from the null predictions (p<0.01). See Materials and Methods and Results for details.

Hence, both cytoachitectonic similarity and distance are significant predictors of presence or absence of ipsilateral and contralateral connections and bivariate models (with both predictors) are better than univariate ones (consisting of each predictor separately). The contribution of each factor is different for ipsilateral and contralateral connections, with the contribution of distance greatly diminishing for the contralateral connections while the contribution of cytoachitectonic similarity remains stable. Moreover, the contribution of each predictor is different for contralateral and ipsilateral connections, with distance, when compared to cytoachitectonic similarity, appearing more tightly related to the presence or absence of ipsilateral connections, while the opposite holds for the contralateral connections (Table 1, Fig. 5).

### Control analyses

The results for the control dataset (Zingg et al. 2014) were in line with the above results on ipsilateral connectivity (Supplementary Table 1). The bivariate model was significantly better than the univariate models (Likelihood ratio test: 50.7 11.2 (p<0.001) when comparing the log likelihood of the bivariate model and the model built only on cytoachitectonic similarity and distance respectively). The prediction analysis for the control dataset demonstrated that significant predictions of connectivity are achieved when using only distance (AUC=0.66), only cytoachitectonic similarity (AUC=0.59) or both (AUC=0.68) (all AUCs p<0.01, permutation tests). Predictions based on distance were significantly better than predictions based on cytoachitectonic similarity and predictions based on both distance and cytoachitectonic similarity were moderately but significantly better than predictions obtained with the use of each predictor separately (p<0.001, permutation tests) (Supplementary Figure 1).

The above results were also significant in all control analyses. First, the control dataset (ipsilateral connections only) confirmed the results of the main analysis (Supplementary Table 1). Second, both datasets underwent a series of “alterations”: omitting two areas from the analysis, adding or deleting 10% of the connections and reassigning cortical types to 10% of the areas. Lastly, a different p-value for determining presence or absence of connections in the main connectivity dataset (Allen Mouse Connectivity Atlas) was used, that is, p<0.01. All control analyses led to results suggesting a significant relation between distance, cytoachitectonic similarity and the presence or absence of cortico-cortical connections. Thus, we are confident that our main results are not severely undermined by methodological limitations in estimating the wiring of mouse cortex (see Limitations and future directions in the Discussion).

Lastly, the addition of a predictor coding for the functional modality similarity of cortical areas did not “explain away” the effect of cytoarchitectonic similarity and distance (Supplementary Table 2). Thus, cytoarchitectonic similarity and distance are fundamental wiring principles, the effect of which is not explained by the functional similarity of cortical areas.

## Discussion

The current results provide novel insights into the cortico-cortical connectional architecture of the mouse brain. We have demonstrated that contralateral connection patterns are significantly similar to the ipsilateral connections and constitute a mirrored but attenuated pattern. Cytoachitectonic similarity and distance relate to the presence or absence of connections, albeit with a different degree of importance for ipsilateral and contralateral connections. In addition, the conjoint usage of these factors leads to more powerful connectivity prediction models when compared to models built with each factor separately. Thus, our results illustrate that cytoachitecture of cortical areas is related to the cortico-cortical connectional architecture (Fig. 6) in addition to the wiring cost principle (e.g. Rubinov et al. 2015).

**Figure 6.**
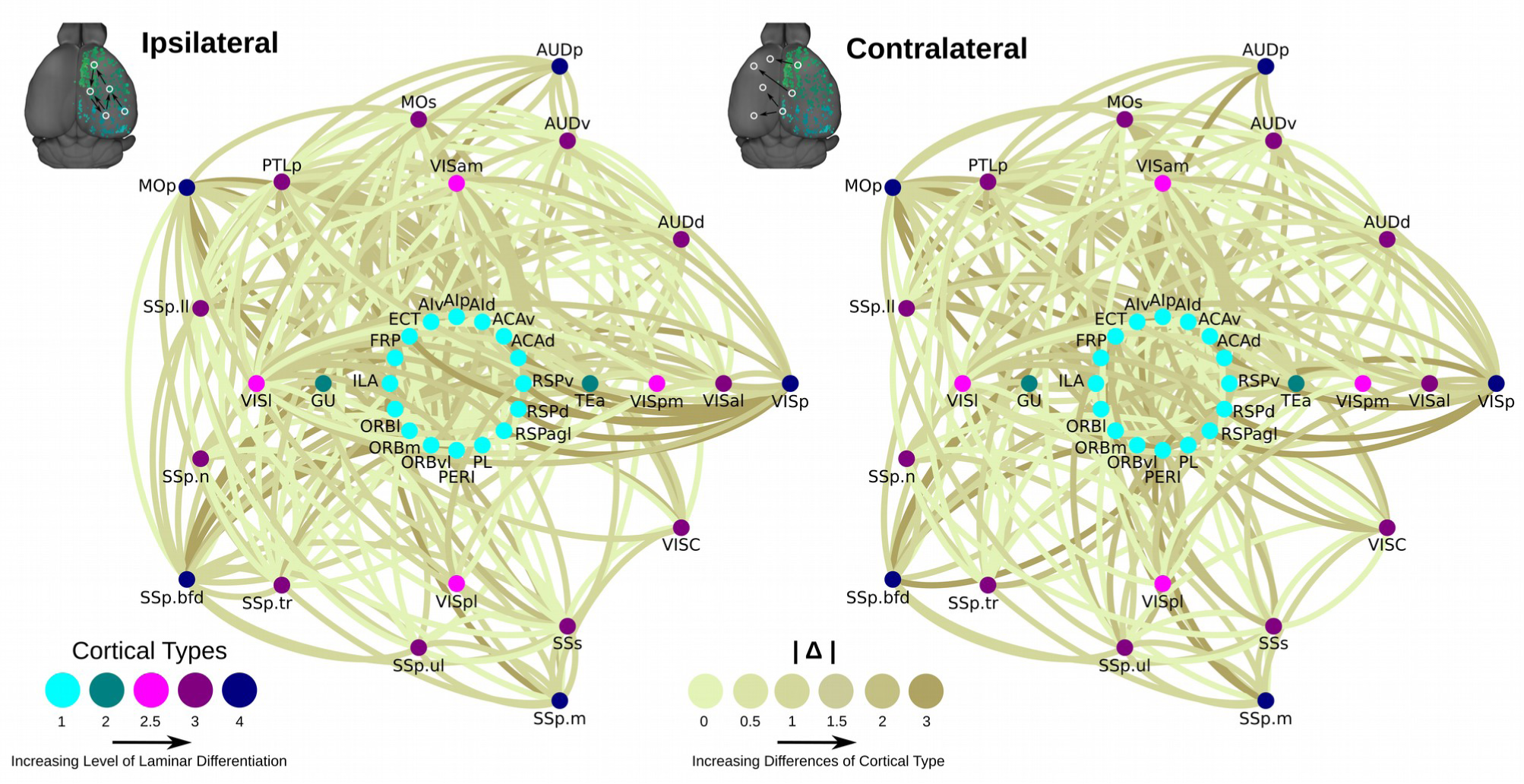
Summary of the mouse cortico-cortical connectional architecture based on cytoarchitectonic similarity. Cortical areas are organized in homocentric circles. The innermost circle contains the less eulaminated areas and successively growing circles correspond to increasingly eulaminated areas. Connections between areas are colour coded based on the absolute difference of their cortical types. Light shades denote similar cortical types and progressively darker shades denote progressively dissimilar cortical types of the interconnected areas. Note that the wiring diagrams are dominated by lighter shades offering a visual summary of the predominant connectivity between areas with similar cortical type. This holds for both ipsilateral and contralateral connectivity.

### Ipsilateral and contralateral connectivity organization

The topology and strength of ipsilateral and contralateral connectivity in the mammalian brain is largely unexamined, especially at the whole brain level. In the macaque prefrontal cortex it was shown that the overall strength of connectivity is much higher for the ipsilateral connections when compared to the contralateral connections (Barbas et al, 2005). Our results suggest that this is also true for the cortico-cortical connections of the mouse at the whole cortex level. Despite an overall higher strength of ipsilateral connections, the patterns for ipsilateral and contralateral connections were moderately to strongly correlated. These findings indicate a mirrored but attenuated connectivity pattern in the mouse, in line with previous observations in the macaque prefrontal cortex (Barbas et al. 2005), suggesting a similar basic topological organization of ipsilateral and contralateral connections in these two species.

The observation that ipsilateral and contralateral profiles of the cortical areas in the mouse do not indicate a segregation of areas in distinct groups (e.g. primary areas versus non-primary areas) (Fig. 3), stands in contrast to findings in the human brain (Stark et al., 2008; Wang et al., 2014). For instance, human cortico-cortical connectivity as estimated in vivo with functional MRI is revealing a distinction between the “primary” and “association” areas, with “association” areas exhibiting prominent ipsilateral connectivity in contrast to “primary” areas (Wang et al., 2014). On the one hand, this finding might suggest a substantial remodeling of ipsilateral and contralateral connectivity in the human brain leading to a more “hemisphere-segregated” architecture. On the other hand, a more prominent ipsilateral connectivity of “association” areas in the human brain, as measured with functional MRI, might emerge due to differences with invasive tract-tracing, that is, the lack of 1:1 correspondence to anatomical connectivity. For instance, structurally unconnected areas can be functionally connected. Lastly, our results demonstrate that homotopic connections are stronger than the average of non-homotopic contralateral connections. This observation is in line with findings in the cat (Lee and Winer, 2008b) and dog (Rajkowska and Kosmal, 1989) cortex. Such findings, obtained from invasive tract-tracing techniques, suggest that the strong homotopic connections observed in the human brain as estimated by functional MRI measurements (e.g. Stark et al., 2008) might also emerge due to strong homotopic anatomical connections.

### Principles of cortico-cortical connectivity

The wiring cost principle already put forward by Ramón y Cajal (1899) has guided recent neuroscientific research focusing on the macroscale connectional architecture of the mammalian brain (Scannell et al., 1995; Chen et al. 2013; Erczsey-Ravasz et al., 2013; Beul et al., 2015a; Beul et al. 2015b). Qualitative observations in the macaque brain suggest that cytoarchitecture of cortical areas is linked to the patterns of cortico-cortical connections (Pandya and Yeterian, 1990). The current results further corroborate and extent such findings by offering important novel insights. First, we demonstrate that distance and cytoarchitectonic similarity are wiring principles also characterizing the mouse brain. This was demonstrated quantitatively by employing two independent datasets that constitute the current best estimate of the macroscale cortico-cortical connectivity of the mouse. Such results resonate well with previous finding in the cat and macaque cortex (Beul et al., 2015a; Beul et al 2105b), highlighting distance and cytoarchitectonic similarity as mammalian-general wiring principles. Second, there is a different degree of association of these principles with the presence or absence of ipsilateral and contralateral connections. The contribution of distance greatly diminishes for the contralateral connections while the contribution of cytoarchitectonic similarity remains stable. Moreover, distance, when compared to cytoarchitectonic similarity, appears more tightly linked to the presence or absence of ipsilateral connections, while the opposite holds for the contralateral connections. This demonstrates that contralateral connections are poorly explained by the wiring cost principle. Hence, these results enrich our understanding of the factors related to the macroscale connectional architecture and extend the set of wiring principles of the mammalian cortex beyond the well documented principle of wiring cost conservation.

### Putative neurobiological mechanisms underlying the observed wiring principles

We have currently demonstrated wiring principles of the mouse cortex. However, principles do not constitute neurobiological mechanistic explanations but offer a quantitative “anchoring point” for further investigating how the observed relations, that is, relations between presence or absence of connections, distance and cytoarchitectonic similarity, come about. With respect to the length of connections, such wiring economy might arise due to a random axonal growth process previously demonstrated in computational modeling giving rise to realistic distributions of physical length of connections (Kaiser et al. 2009). Hence, connections are more likely to be established between areas that are spatially close compared to remote areas. Cytoarchitectonic similarity of areas of the adult brain might reflect similarities in the time window of development during neurogenesis (e.g. Rakic 2002; Charvet et al. 2015). Gradients of neurogenesis suggesting such distinct time windows of neurons populating the distinct cortical areas are documented in the mouse cortex (Smart 1984). Thus, similar time windows in the ontogeny of areas might bias the connections under development to “prefer” areas that will exhibit similar cytoarchitecture in the adult brain, since such areas host neurons that constitute the available origin and more probable projection targets for establishment of connections (for a schematic depiction of this scenario see Fig. 7). Such preferential connectivity is observed in *C.elegans* (Varier and Kaiser 2011), and to a certain extend in the Drosophila connectome (Chiang et al. 2011). While the sketched neurobiological mechanisms are possible scenarios, the current results, in conjunction with similar findings in the cat and macaque cortex, point at a consistent relation between the physical, cytoarchitectonic and connectional brain architecture. Thus, a potentially evolutionary conserved neurobiological mechanism must lie at the heart of such prominent and systematic relation.

**Figure 7.**
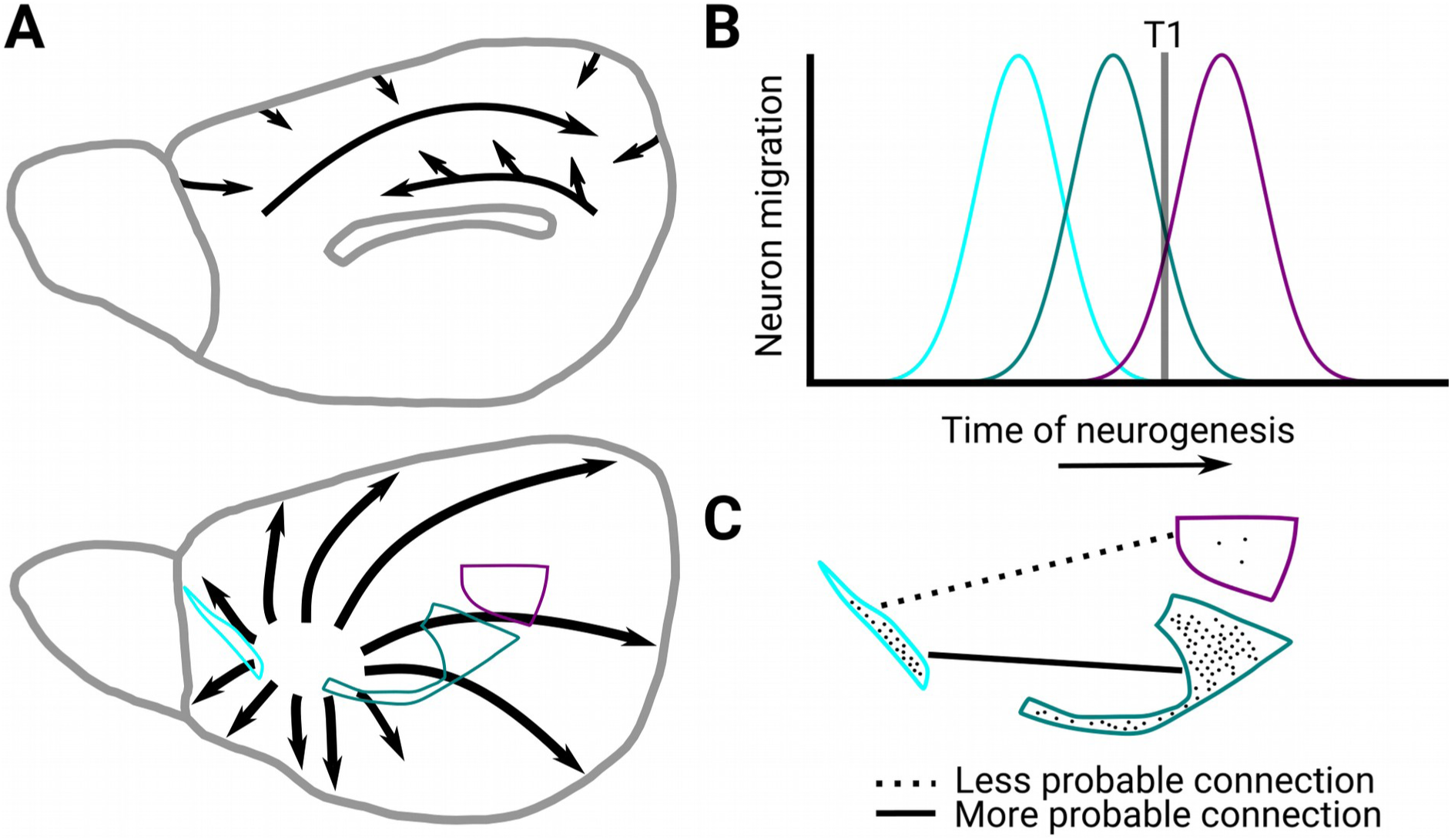
Diagrams illustrating the scenario that the cytoarchitecture of cortical areas in the adult brain reflect distinct time windows of neurogenesis during development resulting in the observed “similar prefers similar” cytoarchitectonic wiring principle. **A**. Gradients of neurogenesis in the mouse cortex. The root of the gradient of neurogenesis in the lateral surface is located near the insula. Gradients in the medial surface exist have two roots. One near the hippocampus and one in the rostral part of the medial surface. Arrows depict the spatial patterns of neurogenesis. These gradients of neurogenesis are based on data presented in (Smart 1984). **B**. Assumed overlapping time windows of neurogenesis for three cortical areas with different cortical types. The less eulaminated area has the earliest onset of neurogenesis while the more eulaminated area has the latest onset of neurogenesis. These differences might arise due to the different position of cortical areas in the gradients of histogenesis depicted in panel A. The curves schematically depict the onset, pick and decline of neurogenesis in the cortical areas. **C**. Establishment of connections at timepoint T1 indicated in panel B might be more plausible between areas that contain neurons generated in similar time windows since they offer more potential “connection partners”.

### Functional considerations

The pattern of presence and absence of connections between cortical areas results in topological network configurations that appear crucial for efficient brain function (e.g. Sporns et al. 2000; Müller-Linow et al. 2008; Moretti and Muñoz 2013). We have illustrated principles that govern such wiring and also putative neurobiological mechanisms shaping this relation. Hence, miswiring leading to pathological brain dynamics and function may be conceived of as a failure to sculpt the cortico-cortical landscape based on such principles. Notably, distance and cytoarchitectonic similarity are also tightly related to connection features like strength (Hilgetag and Grant 2010) and laminar patterns (Barbas 1986; Hilgetag and Grant 2010; Beul et al. 2015a). Strength of connections might lead to a differential functional impact (Vanduffel et al. 1997) and laminar patterns appear important for shaping the spectral channels used for exerted influence between cortical areas (Bastos et al. 2015). Hence, physical distance and cytoarchitectonic similarity relate to many features of cortical connections that in turn are closely linked to distinct functional features. Consequently, the current wiring principles constitute a good guiding thread for uncovering and understanding the links between the distinct levels of cortical organization.

### Limitations and future directions

Certain limitations of our study should be noted. First, the connectivity matrices of (Oh et al. 2014) were derived from a constrained optimization with assumptions such as homogeneity of areas. An assumption of strict area homogeneity might not be neuroanatomically realistic since multiple injections in different parts of what is considered as one cortical area might exhibit different connectivity patterns (e.g. Luppino et al. 2003). Moreover, the automated pipelines used in (Oh et al. 2014) can lead to spurious connectivity estimates due to misregistration and the inability to clearly distinguish fibers of passage from axon terminal. Importantly, our main results were reproduced in an independent dataset (Zingg et al. 2014) that was derived from manual expert annotation and thus is not affected by the above methodological limitations. In addition, the results were unaffected by a set of control analyses. Hence, our results are likely not undermined by a potential update on the status of specific connections of specific areas, such as presence of contralateral connections for some areas that currently were considered absent. Second, Euclidean distance was used as a proxy of wiring cost. Since we examine presence or absence of connections, information on the possible length of non-existent cortical pathways is not available and easy to estimate. Thus, Euclidean distance constitutes a pragmatic estimate of wiring cost for our pertinent research questions.

We have currently examined the relation of distance and cytoarchitectonic similarity based on prior work in the cat (Beul et al. 2015a) and macaque (Beul et al. 2015b) cortex. Previous studies demonstrate a relationship between connectivity and gene expression (French and Pavlidis 2011; Ji et al. 2014) and topological similarity (Costa et al. 2007). Future predictive models could incorporate such information leading to more powerful models explaining the cortico-cortical connectional architecture.

### Conclusions

We examined the topological relation of ipsilateral and contralateral cortico-cortical connectivity in the mouse and the degree to which distance and cytoarchitectonic similarity relate to the presence or absence of connections. Remarkably, despite the striking differences of mouse, cat and macaque brains across space (brain size) and time (phylogeny), a common set of principles appear to underlie their macroscale wiring.

## Acknowledgements

This work was supported by an Alexander von Humboldt fellowship to AG and funding by the German Research Council DFG to CCH (SFB 936/A1). The authors declare no competing financial interests.

